# Neural activity drives directional subarachnoid cerebrospinal fluid flow in the human brain

**DOI:** 10.64898/2026.05.28.728545

**Authors:** Fuyixue Wang, Jonathan R. Polimeni, Lawrence L. Wald, Bruce R. Rosen, Laura D. Lewis, Zijing Dong

**Affiliations:** Athinoula A. Martinos Center for Biomedical Imaging, Massachusetts General Hospital, Charlestown, MA, USA; Department of Radiology, Harvard Medical School, Boston, MA, USA; Richard M. Lucas Center for Imaging, Department of Radiology, Stanford University, Stanford, CA, USA; Department of Electrical Engineering and Computer Science, Massachusetts Institute of Technology, Cambridge, MA, USA

## Abstract

Neural activity is a potential driver of cerebrospinal fluid (CSF) dynamics and brain waste clearance, yet how neural activity shapes localized CSF flow in the subarachnoid space remains poorly understood in humans. Here we introduce an MRI framework that enables concurrent mapping of ultra-slow subarachnoid CSF flow at velocity scales of tens of micrometers-per-second together with hemodynamic activity measured by simultaneous functional MRI in the human brain. Using this framework, we found that neural activity drives spatiotemporally coherent, directional CSF flow in the subarachnoid space near sites of neural activation in awake humans. These flow responses were highly spatially specific, shaped by local cortical and subarachnoid anatomy, while exhibiting oscillatory temporal dynamics modulated by the stimulus cycle and strongly coupled to hemodynamic responses. These findings establish a link between neural activity, vascular dynamics, and localized subarachnoid CSF flow, and provide a framework for investigating neural-activity-driven CSF transport in humans.

## Main

Efficient clearance of metabolic waste is essential for maintaining brain homeostasis and preventing excessive accumulation of neurotoxic byproducts that are associated with neurodegeneration^1-6^. Cerebrospinal fluid (CSF) flow plays a central role in brain clearance by facilitating the transport of solutes through the subarachnoid and perivascular spaces, as described within the framework of glymphatic system^1,3,6,7^. This process is thought to be facilitated by bulk fluid movement within these spaces, which can be driven by physiological factors such as arterial pulsation^8,9^ and respiration^10,11^. Recently, emerging evidence has identified neural activity as a critical driver of CSF dynamics through its coupling with vascular or metabolic processes^12-18^. In particular, human studies combining electroencephalography (EEG) and functional MRI (fMRI) have shown that large-scale neural events, such as slow-wave activity during sleep, accompanied by coordinated hemodynamic changes, induce bulk CSF flow in the 4^th^ ventricle^12^. More recent work has extended these observations to awake states, indicating that task-evoked neural activity can also modulate CSF movement in the 4^th^ ventricle^19^ and increase CSF mobility in the visual cortex^20^. Experimental evidence from animal studies suggests that coordinated neuronal activity generates large-scale ionic waves that regulate CSF–interstitial fluid exchange and drive brain waste clearance^16^, and visual stimulation-induced vasomotion could also enhance clearance in visual cortex of awake mouse^14^. These findings support a model in which neural activity, via its influence on vascular dynamics or metabolic processes, contributes to CSF transport and brain waste clearance. However, beyond the flow observed in the 4^th^ ventricle^21-24^, the neural-activity-driven CSF flow in the subarachnoid space, which is the central hub for brain-wide CSF circulation and transport, remains poorly understood in the human brain, including its spatiotemporal and directional characteristics and coupling to neural-activity-associated vascular responses. This gap stems from the lack of noninvasive methods capable of quantifying ultra-slow subarachnoid CSF dynamics and resolving complex directional flow patterns in the subarachnoid space while simultaneously mapping brain activity.

Here, we introduce an MRI framework that enables simultaneous mapping of subarachnoid CSF flow and neural activity-induced hemodynamic activity in the human brain. It builds on our recent slow-flow-sensitized phase-contrast imaging (SOPHI) acquisition^25,26^, and extends it to allow simultaneous Blood-Oxygenation-Level-Dependent (BOLD) fMRI acquisition leveraging the multi-echo Echo-Planar Time-resolved Imaging (EPTI) readout^27,28^, together with a new CSF flow analysis framework for investigating neural-activity-driven CSF dynamics. This approach enables direct quantification of directional subarachnoid CSF flow at microscopic scales (e.g., <100 μm/s) together with hemodynamic activity and their spatiotemporal coupling. Using SOPHI–EPTI, we revealed spatiotemporally coherent, directional CSF flow in the local subarachnoid space driven by neural activity in the awake human brain, with flow velocities on the order of ∼20–40 μm/s. These directional flow responses were highly spatially specific and shaped by the local subarachnoid anatomy, while exhibiting oscillatory temporal dynamics modulated by the stimulus cycle and strongly coupled to BOLD hemodynamic responses. The observed spatiotemporal flow patterns were robust and reproducible across subjects and repeated scans, with inter-individual variations in flow response amplitude and directionality reflecting differences in individual cortical and subarachnoid structures, indicating a close association between regional brain structure and fluid dynamics. Together, our results reveal that neural-activity-induced hemodynamics can drive spatially organized, directional subarachnoid CSF flow in the human brain with high intra-subject repeatability and inter-subject consistency, establishing technical and biological foundations for investigating the organization of neural-activity-driven CSF flow in humans.

## RESULTS

### Simultaneous slow CSF flow and BOLD fMRI mapping using SOPHI–EPTI

To investigate neural-activity-driven CSF flow in the subarachnoid space, we developed a simultaneous slow CSF flow and BOLD fMRI imaging approach using SOPHI–EPTI, including both acquisition and analysis framework (**Fig. 1**), allowing direct assessment of their spatiotemporal coupling. During acquisition, CSF flow is measured using slow-flow-sensitized phase contrast that provides high sensitivity and specificity to ultra-slow flow at microscopic scales, while hemodynamic BOLD signals are derived from multi-echo magnitude-valued images resolved within a single readout. Specifically, for CSF flow measurement, SOPHI employs a pulsed-gradient spin-echo (PGSE) acquisition with long velocity-encoding times to enhance slow-flow sensitivity, while long echo-times (TEs), spin-echo contrast, and single-shot, distortion-free readout improve CSF specificity by minimizing contamination from blood signals, partial-volume effects, and other confounding factors including physiological noise-induced phase fluctuations and dynamic distortions. Central spin-echoes resolved near the spin refocusing at the center of the EPTI readout—whose contrast is intrinsically less sensitive to confounding phase fluctuations—are extracted and incorporated into a tailored phase-valued data processing pipeline to further suppress confounding background phase. For CSF flow analysis, two approaches are proposed. In the first approach, flow velocities along the three velocity-encoding directions are combined to generate quantitative velocity maps and directional flow vector fields for each task-locked dynamic frame, enabling direct characterization of the spatiotemporal organization of neural-activity-induced CSF flow patterns. In the second approach, we propose a generalized linear model (GLM)-based flow analysis framework applied to the velocity time-series data along each velocity-encoding direction using temporal regressors modeling predicted CSF flow responses, such as the temporal derivative dBOLD/d*t* time-series derived from the predicted BOLD hemodynamic signal, as identified and described in later sections. The resulting t-score maps are generated for all three velocity-encoding directions, capturing both the strength of CSF flow responses (t-score value) and their directional dynamics (t-score sign relative to the temporal regressors along each velocity-encoding direction). This approach enables specific detection of neural-activity-driven CSF flow components coupled to the predicted neural/hemodynamic dynamics instead of unrelated physiological fluctuations, while preserving direction-specific information. For magnitude-based BOLD measurements, since spin-echo acquisitions provide predominantly T_2_-weighted contrast with limited BOLD sensitivity, we resolve multi-echo asymmetric spin-echo images within the same readout using single-shot EPTI, enabling acquisition of T_2_*-weighted contrast^29^ with high sensitivity to both macrovascular and microvascular hemodynamic signals. Standard fMRI analysis is then applied to obtain fMRI activation metrics.

**Fig. 1.**
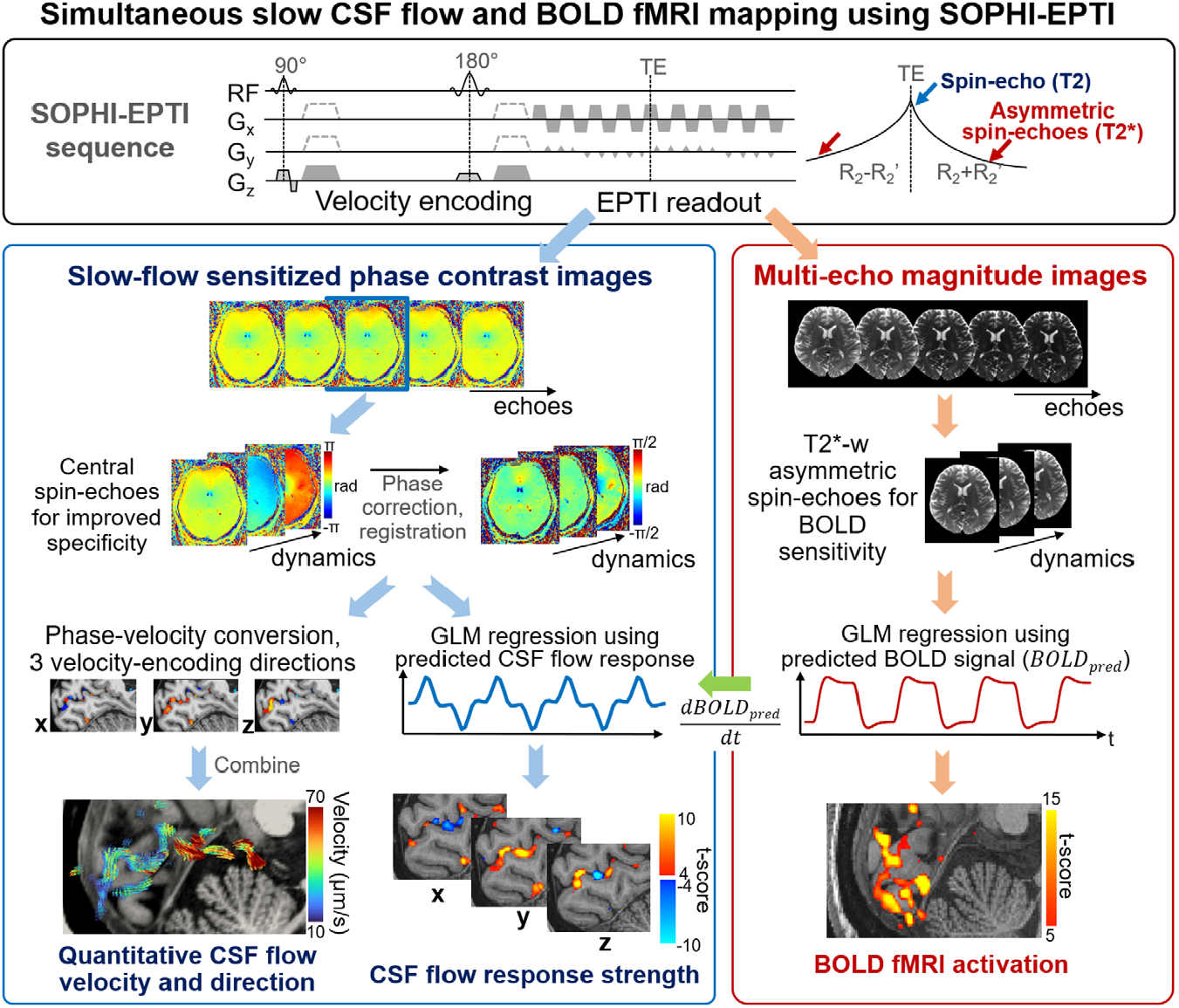
Overview of simultaneous slow CSF flow and BOLD fMRI mapping using SOPHI-EPTI. Top panel, SOPHI–EPTI pulse sequence diagram designed for simultaneous phase-contrast mapping of slow CSF flow and magnitude-based BOLD fMRI. A slow-flow-sensitized sequence with spin-echo-based velocity encoding was employed with a single-shot EPTI readout, enabling acquisition of multi-echo, distortion-free images. Bottom left panel, phase-contrast images from the central spin-echoes are processed with a tailored pipeline with high sensitivity and specificity to CSF signals. Two analysis approaches are then performed for CSF flow data: one generates quantitative flow velocity and directional flow vector field maps for each dynamic frame, whereas the other is a time-series GLM-based analysis to detect neural-activity-driven CSF flow coupled to predicted neural/hemodynamic dynamics, generating flow activation maps with directional dynamic information for each encoding direction. Bottom right panel, magnitude-valued images with T_2_* weighting from asymmetric spin-echoes provide high sensitivity for functional BOLD imaging.

Using SOPHI–EPTI, we performed simultaneous slow CSF flow and fMRI measurements using a 7 Tesla MRI scanner in 11 healthy participants (31 ± 7 years; 8 females, 3 males). Slow CSF flow data were acquired with three orthogonal velocity-encoding directions (posterior–anterior: P–A, inferior–superior: I–S, and left–right: L–R), to enable flow analysis along all directions and 3D flow vector field visualization. Visual stimulation (flickering checkerboard) was applied to evoke neural activity and assess CSF flow responses and their relationship to hemodynamic BOLD responses.

### Visual stimulation evokes spatiotemporally organized oscillatory CSF flow patterns in the subarachnoid space around visual cortex

We first examined CSF flow velocity maps along three orthogonal velocity-encoding directions (**Fig. 2a**) across a full visual task cycle (visual task-locked data with 20 binned time frames; 30 s ON, 30 s OFF). CSF flow responses were observed within the subarachnoid space surrounding the primary visual cortex, exhibiting an oscillatory pattern throughout the task cycle, with elevated velocities occurring during task-state transitions, including stimulus onset (**Fig. 2a** first column, frame 1) and offset (**Fig. 2a** third column, frame 11). In addition, opposite flow directions were observed during stimulus onset and offset across all three velocity-encoding directions. For example, strong forward and upward flow occurred during stimulus onset, whereas backward and downward flow was observed at offset. Task-locked time-series data from representative voxels further illustrate the directional flow responses (**Fig. 2b**), with opposite flow directionality observed during stimulus onset and offset transitions. Notably, voxel-**i** showed stimulus-coupled flow changes along both P–A and I–S directions, whereas voxel-**ii** exhibited predominantly I–S flow modulation. These differences are consistent with local anatomical geometry and curvature of the subarachnoid space, as the former voxel is located within and along an obliquely oriented subarachnoid channel (yellow arrows in **Fig. 2a**), whereas the latter resides in a more vertically oriented channel running along the I–S axis. To further visualize these patterns, we reconstructed three-dimensional CSF flow vector fields around the primary visual cortex (**Fig. 2c**), revealing spatially coherent and organized CSF flow that follows the curvature of the subarachnoid space and exhibits oscillatory dynamics throughout the visual task cycle. Consistently, transition periods immediately following stimulus onset (frame 1) and offset (frame 11) exhibited elevated flow velocities relative to the more stable periods (frame 6 and 16), as reflected by the color coding, together with opposite flow directions, as indicated by the vector orientations. A more direct visualization of the task-induced flow velocities and flow vector fields are shown in **Supplementary Videos 1–3**.

**Fig. 2.**
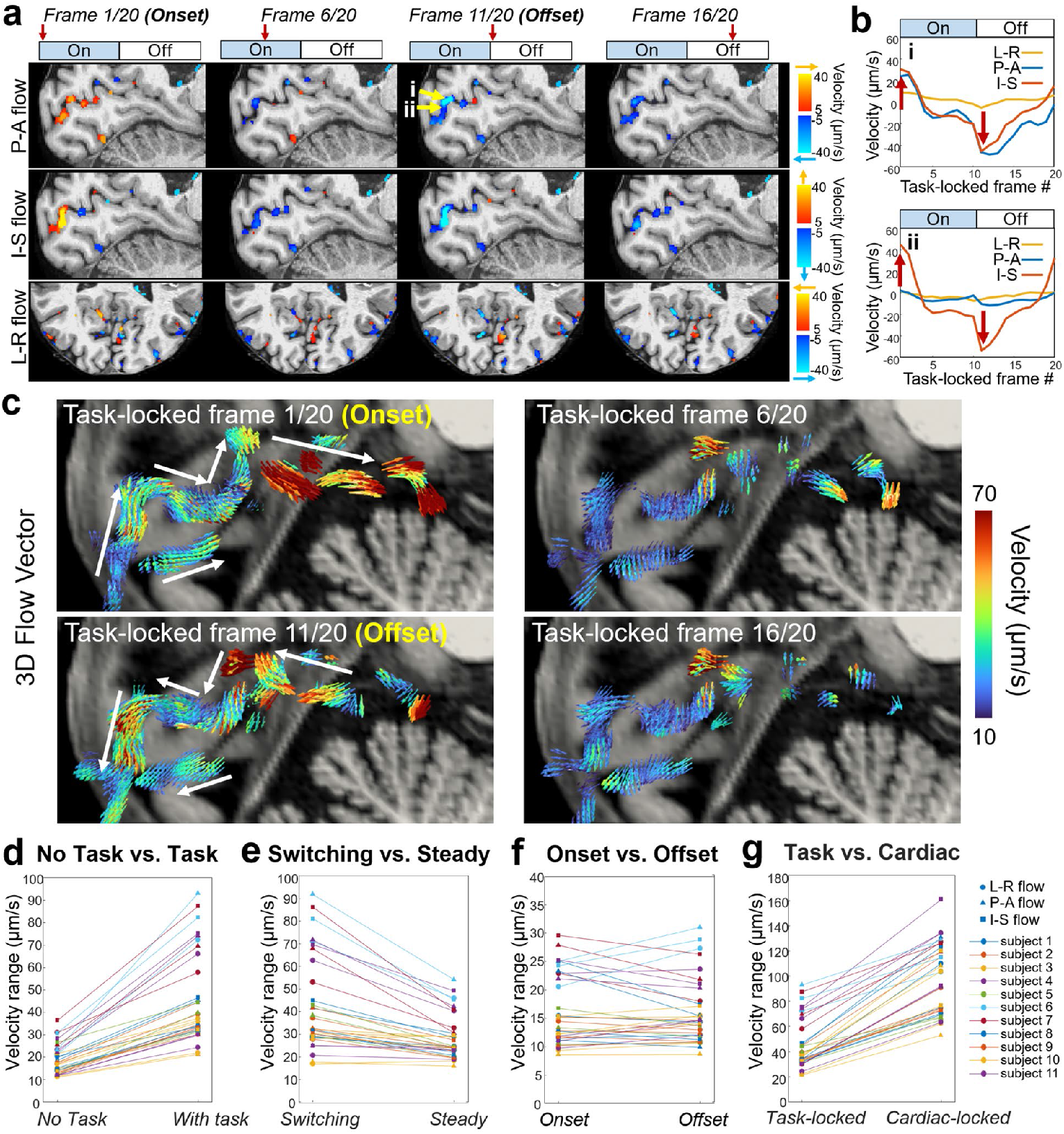
Spatiotemporal subarachnoid CSF flow responses to visual task. **a**, Visual-task locked CSF flow velocity maps along three directions at representative time frames across a visual task cycle. It shows oscillatory changes of the flow velocity and directionality coupled to the stimulus, with elevated velocities and opposite flow directions during stimulus onset (Frame 1/20) and offset (Frame 11/20). **b**, Example task-locked time-series of the 3-directional CSF flow velocities from two representative voxels as indicated by the yellow arrows in **a**. Red arrows indicate the peak velocities during task-state transitions. **c**, 3D flow vector field maps of task-locked CSF flow around primary visual cortex, with white arrows indicating flow directions that follow the anatomical curvature of the subarachnoid space and exhibit opposite flow directions between stimulus onset and offset. Flow vectors are color-coded by velocity amplitude, with vector orientation indicating flow direction and vector lengths normalized. **d**, Comparison between resting-state and visual-task velocity ranges (task-locked). **e**, Comparison between velocity ranges during stimulus-transition periods (frames 1–5 and 11–15) and steady-state periods (frames 6–10 and 16–20). **f**, Comparison between velocity ranges during stimulus onset (frames 1–5) and offset (frames 11–15). **g**, Comparison between task-locked and cardiac-locked velocity ranges. **d–g**, all comparison were performed along the 3 velocity-encoding directions across 11 healthy participants (shapes denote flow directions, and colors denote individual participants). The velocity ranges were averaged within the V1 subarachnoid space.

We then quantified task-driven CSF flow changes by calculating the velocity range (maximum minus minimum velocity across frames of interest) as a measure of flow response amplitude. Within the subarachnoid space surrounding V1, task-evoked CSF flow exhibited substantially larger task-locked velocity ranges compared to resting-state data acquired from the same participants (45.5 ± 20.2 μm/s vs. 18.6 ± 6.6 μm/s; *p* < 0.001; **Fig. 2d**). Further analysis within the task cycle revealed that transition periods immediately following stimulus onset and offset (frames 1–5 and 11–15) exhibited significantly greater velocity ranges than steady-state periods (frames 6–10 and 16–20) (42.3 ± 20.8 μm/s vs. 28.7 ± 9.8 μm/s; *p* < 0.001; **Fig. 2e**), suggesting that CSF flow is primarily enhanced during task-state transitions. No significant difference was observed between onset and offset transition periods (**Fig. 2f**), suggesting a relatively symmetric flow modulation across the task cycle. We next compared task-locked CSF flow with cardiac-locked flow derived from retrospective cardiac gating in the same voxels of V1 subarachnoid space. Cardiac-locked velocity ranges exhibited greater amplitudes than task-locked flow (96.9 ± 28.0 μm/s vs. 45.5 ± 20.2 μm/s; *p* < 0.001; **Fig. 2g**), consistent with prior understanding that arterial pulsation is a dominant driver of CSF dynamics^8,9,25,26^. However, task-evoked CSF flow reached approximately ∼40–50% of the cardiac-driven amplitude, demonstrating that neural activity contributes substantially and measurably to subarachnoid CSF flow in the human brain.

### Neural-activity-evoked directional subarachnoid CSF flow is temporally coupled to dBOLD/d*t* dynamics

A hypothesized mechanism underlying neural-activity-driven CSF flow involves the hemodynamic response and its associated changes in cerebral blood volume (CBV)^12^ (**Fig. 3a**). Periods of stimulus-evoked neural activity exceeding ∼10 s can induce a localized hemodynamic response characterized by arterial/arteriolar dilation followed by venular/venous expansion^30-32^, generating pressure changes in the surrounding subarachnoid space^33-35^ that can drive directional CSF flow. To test this hypothesized model, we analyzed the simultaneously acquired slow CSF flow and BOLD fMRI data during visual stimulation. Strong BOLD activation was observed in the visual cortex (**Fig. 3b**, top panel) with BOLD signal time series following the task paradigm as expected (**Fig. 3c**, top panel). CSF flow responses were observed within the surrounding subarachnoid space (**Fig. 3b**, bottom panel) near sites of neural activation. In contrast to BOLD, they exhibited distinct temporal features (**Fig. 3c**, middle panel) characterized by elevated velocities during task-state transitions (± 30 μm/s), with flow velocities ramping toward peak values before returning toward baseline during steady-state periods, and with opposite flow directions and ramping dynamics during stimulus onset and offset. The overall CSF flow time-series closely resembled the temporal derivative of the BOLD signal (dBOLD/d*t*) (**Fig. 3c**, bottom panel), as demonstrated in an example CSF voxel around V1 (*r* = 0.90, *p* < 0.001). This strong temporal correlation between CSF flow response and dBOLD/d*t* is consistent with the hypothesized model in which CBV changes occur predominately during task-state transitions, with opposite CBV changes associated with vascular expansion and relaxation at stimulus onset and offset respectively. Note that the CSF flow time-series was averaged across multiple runs and temporally smoothed by a median filter (window size = 3), so the effects from other physiological drivers such as cardiac pulsation are diminished, which would otherwise introduce additional fluctuations (manifesting as noise) into the time-series data.

**Fig. 3.**
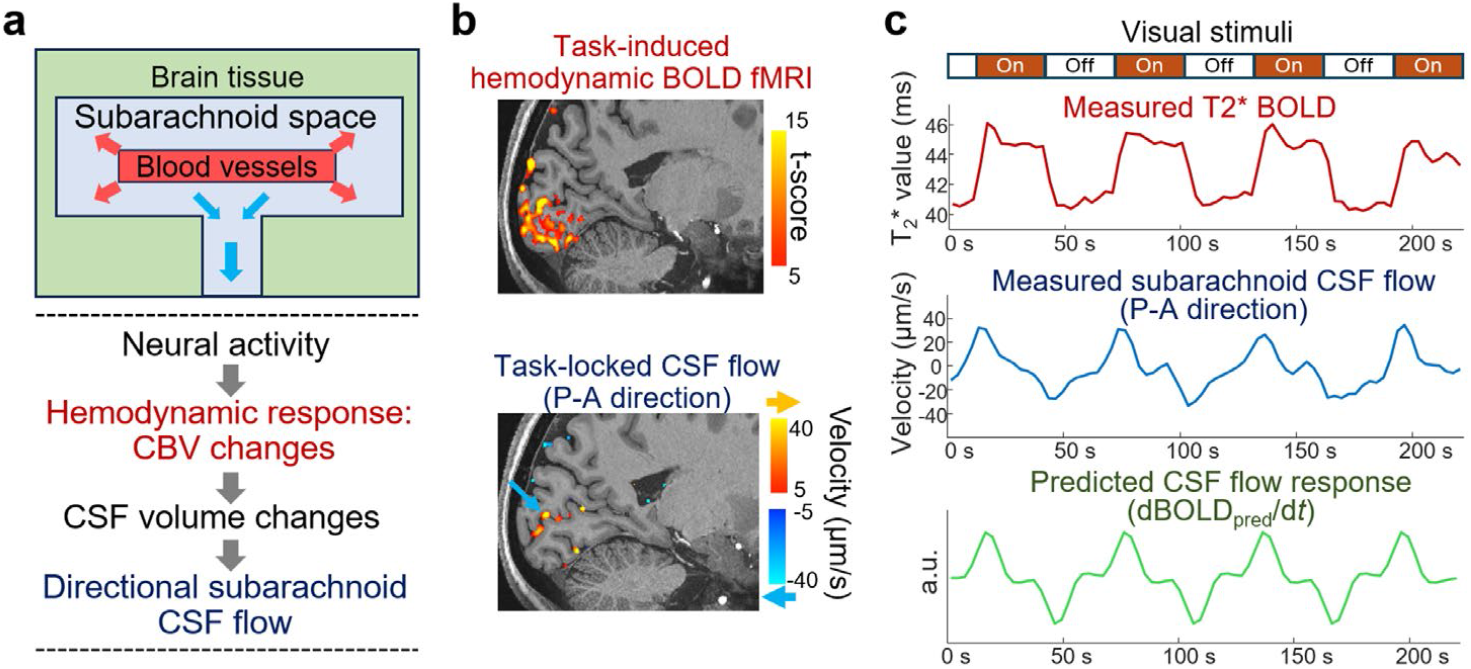
Coupling between subarachnoid CSF flow and hemodynamic responses. **a**, Hypothesized model of neural-activity-driven subarachnoid CSF flow mediated by hemodynamic CBV changes. **b**, Visual task-induced BOLD fMRI activation and P–A directed CSF flow velocity maps at a representative time frame (during stimulus onset), showing strong hemodynamic and CSF flow responses around the primary visual cortex. **c**, Top panel, BOLD response averaged within the V1 ROI; middle panel, measured subarachnoid CSF flow from an example voxel (blue arrow in **b**); bottom panel, the predicted CSF flow response (dBOLD_pred_/d*t*) based on the hypothesized model.

### Time-series flow analysis reveals spatially specific and reproducible directional CSF flow responses across subjects and scans

We then employed our General Linear Model (GLM)-based flow analysis framework to the phase-contrast data for specific detection of task-locked CSF flow components coupled to the predicted temporal and directional dynamics. Given the strong temporal coupling between the measured CSF flow and dBOLD/d*t*, we used the temporal derivative of the predicted BOLD signals (dBOLD_pred_/d*t*) as the primary regressor to model CSF flow responses along each velocity-encoding direction (**Fig. 4a**, left panel). The resulting whole-brain t-score maps revealed highly spatially specific CSF flow responses localized to the subarachnoid space surrounding the visual cortex near the sites of neural activation, consistently observed across t-score maps from all three velocity-encoding directions (**Fig. 4a**, right panel for P–A flow and **Extended Data Fig. 1** for all directions), demonstrating the specificity of this approach for detecting task-locked CSF flow responses. The flow t-score maps captured both the strength of the CSF flow response (t-score amplitude) and its directional dynamics for each velocity-encoding direction, with positive and negative values reflecting opposite flow direction dynamics. For example, a positive t-score in the P–A velocity-encoding direction indicates forward (posterior-to-anterior) flow during stimulus onset followed by flow reversal during offset, whereas a negative t-score indicates the opposite. Similarly, a positive t-score in I–S flow reflects upward flow during onset and downward flow during offset.

**Fig. 4.**
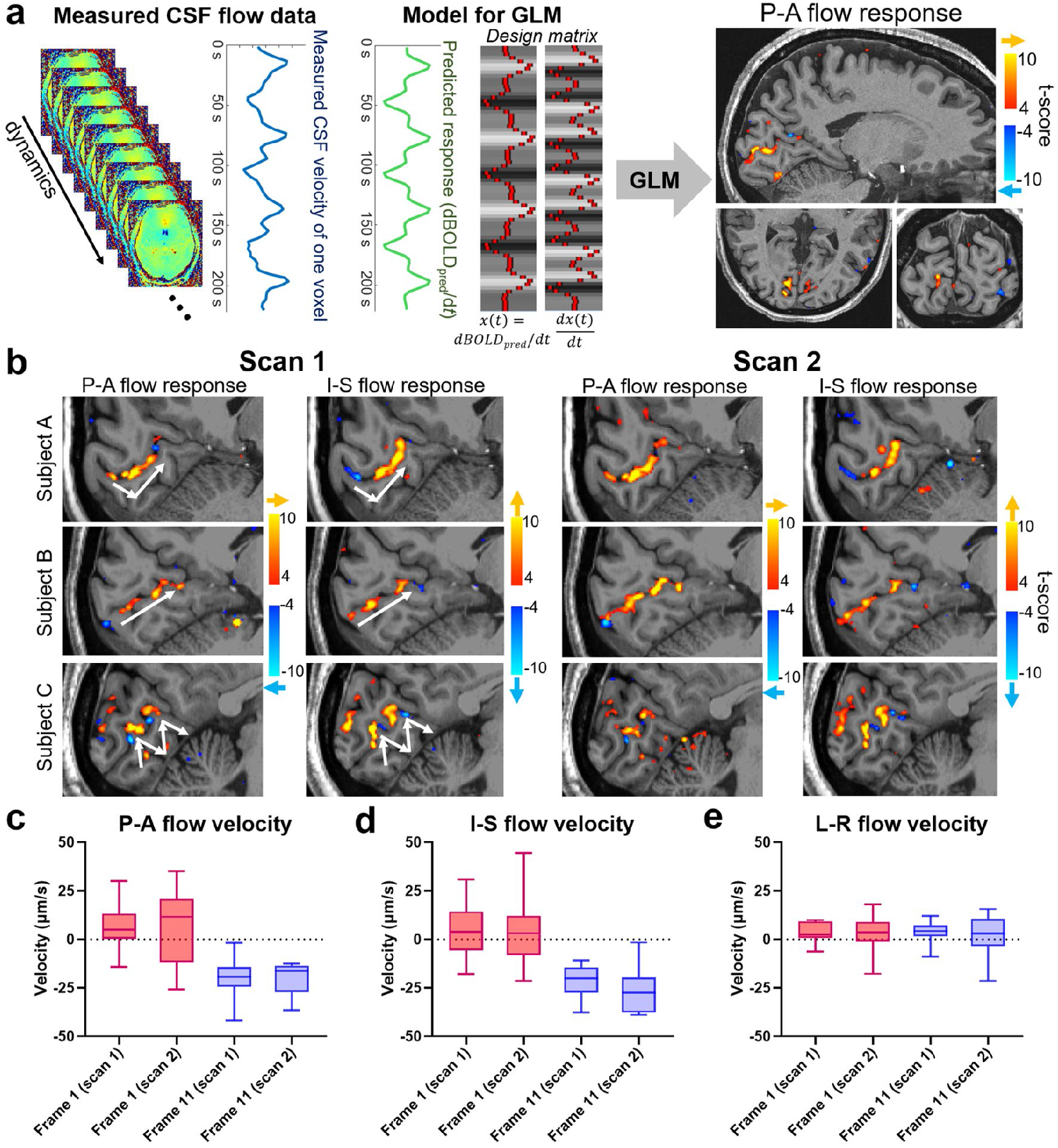
Reproducible task-evoked directional subarachnoid CSF flow responses revealed by GLM-based flow analysis. **a**, Illustration of the GLM-based flow analysis framework. Given the strong temporal coupling between the measured CSF flow and dBOLD/d*t* (left panel), the predicted CSF response was modeled using dBOLD_pred_/d*t* (the temporal derivative of the predicted BOLD response) as the primary regressor (middle panel). The resulting t-score maps (right panel) represent both the strength of the CSF flow response (t-score magnitude, indicated by color) and its directional dynamics for each flow encoding direction, with positive and negative values reflecting opposite flow direction dynamics (colored arrows on the color bar indicate the flow direction during stimulus onset). **b**, T-score maps for P–A and I–S directed flow from three example subjects, each with two repeated scans. White arrows indicate the flow directions shaped by the local curvature of the subarachnoid space anatomy. **c**, Box plots of flow velocities (averaged within the V1 subarachnoid space) across all participants, comparing stimulus onset and offset frames across repeated scans, along three flow-encoding directions. The center line indicates the median, the box denotes the interquartile range, and the whiskers represent the minimum and maximum values.

The resulting t-scores revealed highly reproducible spatiotemporal flow patterns across scan–rescan sessions and across subjects, with inter-individual differences primarily shaped by the local curvature of the individual subarachnoid space. Representative examples are illustrated in **Fig. 4b**: in Subject A, a V-shaped subarachnoid geometry gave rise to forward flow coupled with downward-to-upward transitions along the I–S axis; in Subject B, a relatively straight geometry supported predominantly forward and upward flow; and in Subject C, a more complex M-shaped subarachnoid space resulted in alternating upward and downward flow patterns that follow the underlying anatomy; all patterns were consistent in the repeated scans. Furthermore, analysis across all 11 participants, each with both scan and rescan sessions, examining flow velocities during onset and offset transition frames within the V1 subarachnoid space across three velocity-encoding directions demonstrated that the overall flow modulation was highly consistent across subjects and across scans and re-scans, with anterior–superior directed flow at stimulus onset (**Fig. 4c**,**d**, red bars) and posterior–inferior directed flow at offset (**Fig. 4c**,**d**, blue bars). Although task-driven velocity changes were evident along the L–R direction (**Fig. 2a** and **Extended Data Fig. 1**), the L–R flow velocities in **Fig. 4e** were low due to symmetric cancellation within the ROIs. Together, these results show that visual task-driven CSF flow responses are highly spatially specific to the visual cortex as revealed by the time-series GLM analysis, with spatiotemporal patterns highly reproducible across subjects and scans and shaped by the local subarachnoid space anatomy.

In addition to the reproducibility of the spatiotemporal flow pattern, we further assessed the repeatability for quantifying task-evoked CSF flow and BOLD fMRI measurements in terms of velocity range and t-score across all participants (*N* = 11). For CSF flow, task-locked velocity ranges within V1 demonstrated high repeatability (**Extended Data Fig. 2a**; *r* = 0.99, *p* < 0.001; slope = 1.04, intercept = −2.35), and the CSF flow t-scores also showed strong correlations across sessions (**Extended Data Fig. 2b**; *r* = 0.81, *p* < 0.001; slope = 1.10, intercept = −0.10). Both the velocity range and the t-score maps across the three velocity-encoding directions exhibited highly consistent spatial patterns and magnitudes within V1 (representative examples in **Fig. 4b** and **Extended Data Fig. 2e**). For the simultaneously acquired BOLD fMRI, both t-scores and percent signal changes averaged within the V1 ROI demonstrated high repeatability across sessions (**Extended Data Fig. 2c**; t-score: *r* = 0.96, *p* < 0.001; slope = 0.94, intercept = 1.27; **Extended Data Fig. 2d**; percent signal change: *r* = 0.98, *p* < 0.001; slope = 1.00, intercept = 0.13). Scan–rescan t-score maps also exhibited similar spatial distributions and activation levels (representative examples in **Extended Data Fig. 2f**). Together, these results demonstrate reliable and repeatable CSF flow and BOLD measurements obtained by the proposed framework for investigating neural-activity-driven CSF flow dynamics in the human brain.

### Visual stimulation-induced CSF flow velocity scales with V1 gray matter and subarachnoid CSF volumes

Despite the coherent directionality of task-evoked CSF flow across subjects, we observed substantial inter-individual variability in the magnitude of flow velocity (**Fig. 2e**), with task-locked velocity ranges spanning ∼20–90 μm/s. To identify potential contributors to this variability, we examined relationships between CSF flow magnitude and several structural and physiological factors, including cortical gray matter and CSF volumes as well as BOLD activation strength. Notably, task-evoked CSF flow velocity range showed significant positive correlations with both V1 cortical gray matter volume (**Fig. 5a**; *r* = 0.78, *p* = 0.0045) and V1 subarachnoid CSF volume (**Fig. 5b**; *r* = 0.75, *p* = 0.0084). These two structural measures were themselves strongly correlated (**Extended Data Fig. 3a**; *r* = 0.78, *p* = 0.0045), likely reflecting their shared dependence on cortical folding and sulcal geometry. Two representative subjects (Subjects 2 and 6, shown in **Fig. 5a–b**) provide an illustrative example of this relationship, with the subject exhibiting larger V1 gray matter volume showing greater task-evoked velocity ranges along both the P–A and I–S directions (**Fig. 5e**). In contrast, no significant relationship was observed between task-evoked CSF flow velocity range and whole-brain gray matter volume (**Extended Data Fig. 3b**; *r* = 0.45, *p* = 0.166), indicating that visual task-evoked CSF flow is specifically linked to local structural features of the visual cortex rather than global brain structure.

**Fig. 5.**
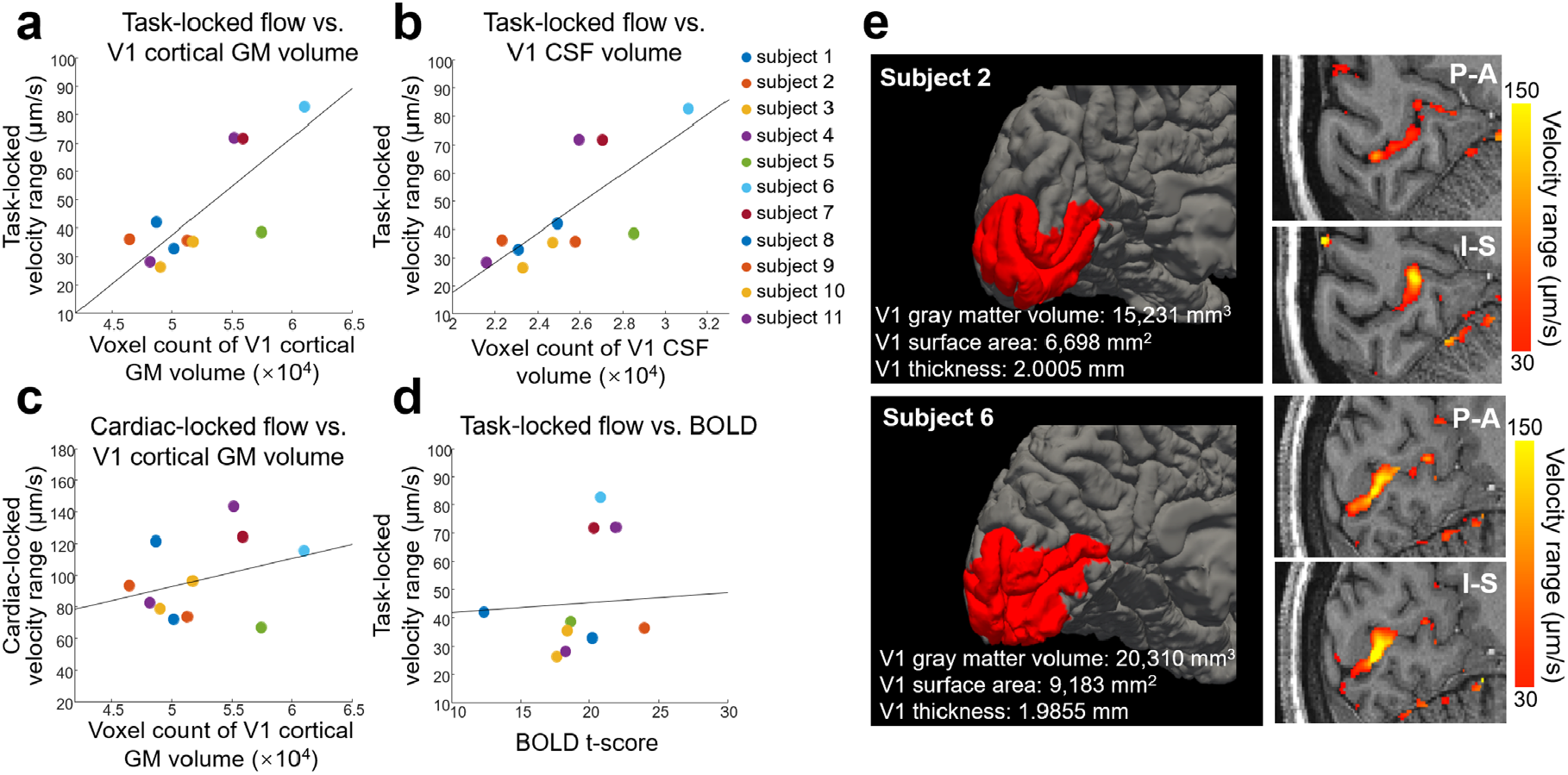
Visual task–evoked CSF flow velocity correlates with V1 cortical gray matter and subarachnoid CSF volumes. **a**, Scatter plot of task-locked velocity range averaged within V1 subarachnoid space versus V1 cortical gray matter volume across all subjects (*N*=11). Individual subjects are denoted by filled circles of different colors. **b**, Scatter plot of task-locked velocity range averaged within V1 subarachnoid space versus V1 subarachnoid CSF volume. **c**, Scatter plot of cardiac-locked velocity range averaged within V1 subarachnoid space versus V1 cortical gray matter volume. **d**, Scatter plot of task-locked velocity range averaged within V1 subarachnoid space versus BOLD t-score averaged within V1. **e**, V1 cortical segmentations from two example subjects with different V1 cortical gray matter volumes, alongside P–A and I–S flow velocity maps showing higher velocities observed in Subject 6 with larger V1 cortical gray matter volume.

Substantial variability in V1 size and surface areas across healthy individuals has been well documented and is associated with differences in visual processing and perception^36,37^. The observed relationship between V1 structure and CSF flow magnitude suggests an important role of regional anatomical organization in shaping neural-activity-driven fluid dynamics. In addition, we found that cardiac-locked CSF velocity range measured from the same voxels within V1 did not significantly correlate with V1 cortical GM volume (**Fig. 5c**; *r* = 0.32, *p* = 0.34), indicating that neural activity–driven CSF flow is more closely linked to regional cortical and subarachnoid structure than cardiac-driven CSF flow. This distinction may reflect differences in the underlying driving mechanisms, with visual stimulus-evoked hemodynamics producing slower, more localized CSF responses, in contrast to the more rapid and widely propagating cardiac pulsatility-evoked responses. Furthermore, CSF flow magnitude was not significantly correlated with BOLD fMRI metrics across subjects, including t-scores (**Fig. 5d**; *r* = 0.08, *p* = 0.81) and percent signal change (**Extended Data Fig. 3c**; *r* = −0.12, *p* = 0.72).

## Discussion

By introducing a non-invasive MRI framework for simultaneous mapping of ultra-slow CSF flow and hemodynamic activity, our central finding is that neural activity drives spatiotemporally coherent, directional CSF flow in the awake human brain in the subarachnoid space near sites of neural activation, extending previous observations of ventricular CSF flow that reflected a global flow measurement^12,19^, to localized CSF flow responses with detailed spatial organization and regional specificity. We found that neural-activity-driven flow responses exhibited high spatial specificity with localized flow responses whose magnitude and directionality were shaped by local cortical and subarachnoid anatomy. Temporally, the subarachnoid flow responses showed oscillatory dynamics modulated by the stimulus cycle, with elevated velocities occurring during stimulus transition periods and opposite flow directions between stimulus onset and offset. The temporal dynamics of the directional subarachnoid CSF flow were strongly coupled to the hemodynamic responses (e.g., dBOLD/d*t*). Together, these findings reveal a spatially organized and anatomically constrained CSF flow system that is dynamically modulated by neural activity, with prominent flow velocities on the order of tens of micrometers per second.

Our work represents an important technical advance for CSF flow imaging through the development of simultaneous CSF flow and BOLD fMRI acquisition together with a new CSF flow analysis framework. In particular, the proposed acquisition achieves high sensitivity to both ultra-slow flow (e.g., 10–50 μm/s) and BOLD fMRI, capable of measuring complex quantitative, directional flow in three dimensions together with hemodynamic signals across fine spatial and temporal scales. The proposed CSF flow analysis framework includes direct spatiotemporal mapping of the flow velocity and direction, together with a GLM-based analysis established based on the observed temporal characteristics of neural activity-coupled CSF flow, enabling more specific detection of flow components coupled to the predicted responses while suppressing contributions from unrelated physiological fluctuations. The robustness and reproducibility of the resultant flow patterns, directionality dynamics, and quantitative metrics across participants and scan sessions validated the proposed framework. Moreover, the measured flow patterns closely followed the curvature of the subarachnoid space, supporting their physiological origin rather than reflecting artifactual signal contributions. The proposed framework provides a generalizable strategy for investigation of CSF flow responses across a broad range of task and resting-state paradigms, enabling future studies of neural, vascular, and physiological modulation of CSF dynamics.

Our findings suggest a potential new avenue for modulating CSF dynamics—and possibly glymphatic clearance—in localized brain regions, such as subarachnoid space surrounding specific cortical areas, through controlled neural stimulation (e.g., gamma entrainment^38-40^). Our results showed that neural activity constitutes a substantial contributor to subarachnoid CSF dynamics, with task-evoked flow magnitudes near activation sites reaching approximately 40–50% of local cardiac-driven flow— the dominant driver of CSF dynamics. The highly consistent and spatially organized CSF flow patterns observed across subjects and scan-rescans further suggest that neural activity provides a reliable and reproducible mechanism for modulating subarachnoid fluid dynamics. This may be particularly important because disruptions in local CSF flow may contribute to local accumulation of toxic proteins, while local protein accumulation itself may further impair CSF circulation and clearance, potentially creating a worsening feedback cycle. The ability to modulate local fluid dynamics may therefore provide an important strategy for influencing clearance processes. Our findings that the strongest CSF flow modulation occurred during task-state transitions, together with the comparable flow magnitudes observed during stimulus onset and offset, also provide insight into how task paradigms may be designed to more effectively modulate CSF dynamics. The highly localized nature of these responses suggests that distinct neural or behavioral paradigms could be used to selectively modulate CSF dynamics in targeted brain regions.

Through characterization of the flow dynamics, we gained insights into the mechanisms underlying neural-activity-driven CSF flow. The strong temporal coupling between subarachnoid CSF flow and dBOLD/d*t* supported a mechanism in which neural activity-evoked hemodynamic changes, particularly CBV fluctuations, generate local pressure gradients that drive directional CSF motion. The observed direction reversals between stimulus onset and offset further supported this model, reflecting bidirectional vascular expansion and relaxation associated with task-evoked responses. In addition, neural-activity-driven CSF flow was spatially localized and showed strong associations with regional cortical gray matter and subarachnoid structures, suggesting that cortical folding and sulcal geometry may influence both the spatial confinement of vascular responses and the anatomical and biomechanical properties of the surrounding CSF space, thereby shaping both the amplitude and the directionality of the CSF flow response. These characteristics differed from local cardiac-locked CSF flow, which showed no significant correlation with local cortical or subarachnoid CSF volumes. Such differences may reflect distinct underlying driving mechanisms: neural activity-linked CSF flow arises from slower and spatially confined hemodynamic responses, whereas cardiac-driven flow reflects rapid and globally propagating systemic pulsations that are not restricted to local brain regions. Furthermore, despite the close temporal coupling between CSF flow and BOLD dynamics, we observed no significant correlations between the overall amplitude of CSF flow and conventional BOLD metrics (e.g., t-scores and percent signal change), suggesting that the underlying determinants of CSF flow amplitude are not fully reflected by conventional BOLD measurements. This may in part be because BOLD signals are influenced by multiple physiological factors and do not provide a direct or specific measure of CBV^41,42^, or that they predominantly reflect venous hemodynamics rather than arterial-side CBV fluctuations that may play a larger role in driving CSF motion. Another important reason is that BOLD measures are less quantitative and can vary substantially across subjects. Nevertheless, BOLD fMRI remains the predominant imaging contrast for characterizing neural activity and hemodynamic responses because of its unparalleled sensitivity and spatiotemporal resolution, making it particularly valuable for task-based, resting-state, and sleep studies^12,43^.

The ability to non-invasively map neural-activity-driven CSF flow opens several important directions for future research. One promising direction is the investigation of sleep-related CSF dynamics, where large-scale neural oscillations have been linked to enhanced glymphatic clearance^12,18^. Applying this framework could enable spatially resolved mapping of spontaneous neural-activity-associated CSF circulation patterns during sleep, where simultaneous BOLD measurements are particularly critical^12,43^ compared with task-based paradigms. Another key application is in neurological and neurodegenerative diseases, where disruptions in CSF flow and clearance have been implicated in conditions such as Alzheimer’s disease^6,44,45^ and cerebral small vessel disease^46,47^. The ability of SOPHI–EPTI to map both spatial and temporal organizations of CSF flow may enable discovery of novel biomarkers for detecting early region-specific alterations in fluid dynamics and neurovascular coupling, and advance the understanding of disease mechanisms, including through applications to intervention paradigms such as neuromodulation and pharmacological manipulations. While the current study focused on visual stimulation, the proposed framework is generalizable to other brain regions, cognitive tasks, and behavioral states, providing a tool for investigating how neural activity shapes CSF dynamics across diverse functional and physiological conditions.

In summary, we introduced a non-invasive MRI framework for simultaneous mapping of ultra-slow CSF flow and neural-activity-induced hemodynamic responses in the human brain. Using this framework, we demonstrated that neural activity drives spatially organized, directional subarachnoid CSF flow with strong spatiotemporal coupling to hemodynamic dynamics, high intra-subject repeatability, and robust inter-subject consistency. These findings establish both technical and biological foundations for future investigations of neural-activity-driven CSF transport and region-specific flow pathways with rich spatiotemporal detail, ultimately advancing our understanding of human CSF circulation and its role in brain waste clearance and the development of neurological diseases.

## METHODS

### SOPHI–EPTI acquisition for simultaneous CSF flow mapping and BOLD fMRI

As illustrated in **Fig. 1a**, a pulsed-gradient spin-echo (PGSE) sequence with slow-flow-sensitized velocity encoding^26,48,49^ combined with a single-shot EPTI readout^27,28,50^ was employed for data acquisition. EPTI generates multi-echo, distortion-free phase- and magnitude-valued images within each readout (95 echoes in this study), with echoes sampled at a defined echo spacing (ESP) interval. For phase-contrast flow measurements, SOPHI employs an extended velocity-encoding time (compared to that of conventional phase-contrast MRI) to achieve a low velocity-encoding value (VENC) without substantial diffusion-induced CSF signal loss, thereby improving sensitivity to slow flow^25,51,52^. A VENC of 1.6 mm/s was used in this study. SOPHI also employs several additional acquisition strategies to enhance CSF specificity: blood signal was suppressed through the use of a long TE imaging at 7T and further minimized by velocity-encoding-induced dephasing (effective b ≈ 80 s/mm^2^) of fast-moving spins; respiration-related field variations and spatiotemporal phase fluctuations were mitigated by the spin-echo EPTI readout, and only central readout echoes near the spin-echo (28 out of 95 echoes) were used for phase-contrast estimation; the distortion-free EPTI readout also eliminated image geometric distortions and their temporal changes, improving the fidelity of phase-valued time-series measurements, which is particularly critical for resolving CSF flow within the small and confined subarachnoid space. For magnitude-based BOLD measurements, multi-echo images were averaged to increase signal-to-noise ratio (SNR), and the inclusion of asymmetric spin-echo images provided T_2_* contrast to enhance BOLD sensitivity^53,54^. Quantitative T_2_* maps were also fitted from the multi-echo data, providing a more specific measure of BOLD-related T_2_* contrast changes.

### Experimental design and data acquisition

All experiments were performed on a 7T MRI system (MAGNETOM Terra, Siemens Healthineers, Erlangen, Germany) using a 32-channel Nova head coil (Nova Medical, Wilmington, MA, USA). All participants provided written informed consent under an Institutional Review Board (IRB)–approved protocol in accordance with institutional guidelines. Eleven healthy young adults participated in this study (*N* = 11; 31 ± 7 years; 8 females, 3 males). Cardiac physiological signals were recorded during all scans using a fingertip piezoelectric sensor.

Both visual task and resting-state data were acquired using SOPHI–EPTI. Imaging parameters were: 2 mm isotropic resolution; VENC = 1.6 mm/s; FOV = 216 × 216 × 92 mm^3^; TR = 3000 ms; multiband factor = 2; echo spacing = 0.6 ms; and 95 EPTI echoes spaced apart by an echo spacing of 0.6 ms with the spin echo centered at TE = 88 ms. For each velocity-encoding direction, 80 dynamics were acquired in 4 minutes (3 s per image volume), including 6 non-velocity-encoded calibration volumes at the beginning of each run followed by 74 velocity-encoded volumes. To assess task-evoked CSF flow, a block-design visual stimulation paradigm was employed (30 s flickering checkerboard alternating with 30 s gray field, starting with a 30 s baseline fixation period). Two repeated three-directional runs were acquired with the visual task for repeatability evaluation, along with an additional resting-state scan for comparison. Before each three-directional run, a fast EPTI calibration scan lasting ∼50 s was acquired to estimate coil sensitivities and B_0_ field maps for subspace reconstruction. All scans were performed with participants in the supine position. High-resolution anatomical images were acquired using a T_1_-weighted MPRAGE sequence at 0.75 mm isotropic resolution for tissue segmentation.

### Image reconstruction and processing

Raw k-space data were reconstructed using standard EPTI reconstruction with a subspace approach^55,56^ in MATLAB (MathWorks, Natick, MA, USA) and BART^57,58^ to generate multi-echo magnitude and phase images. For phase-contrast CSF flow data, central echoes were averaged to generate a single phase volume for each time frame for input into the phase processing pipeline. During the phase processing pipeline, background phase correction was first performed using third-order spatial polynomial fitting on each slice and time frame, excluding CSF voxels, to estimate and remove spatially smooth phase variations arising from respiration, motion, and eddy currents. This procedure minimally affected CSF flow-induced phase, which is spatially higher in frequency. AFNI^59,60^ was then used to perform rigid motion correction, with motion parameters estimated from all-echo-averaged magnitude volumes and applied to both magnitude- and phase-valued data. Nearest-neighbor interpolation was used to preserve voxel-wise acquisition timing. Phase differences between each flow-encoded volume and the non-velocity-encoded calibration volume were then computed. A second third-order spatial polynomial fitting was applied to remove residual background phase bias. Phase values were then converted to velocity using the formula velocity = (phase/π)×VENC. CSF masking was applied by combining an intensity-based CSF mask (generated by selecting voxels with signal intensities greater than four times the mean signal) with a segmentation-derived CSF mask generated using the FAST tool in FSL^61,62^. Spatial smoothing was performed within the CSF mask for phase-contrast data using a Gaussian kernel with a full-width-at-half-maximum (FWHM) of 3 mm. For CSF flow analysis, flow velocities along the three velocity-encoding directions are combined to compute quantitative velocity maps and directional flow vector fields for each dynamic frame. For GLM-based analysis of neural activity-evoked CSF flow, a temporal median filter (window size = 3) was applied to the flow data to suppress noise and cardiac-related fluctuations; the same processing was applied to the corresponding regressors used in the GLM analysis, such as the predicted CSF flow response—i.e., the dBOLD_pred_/d*t* regressor. The dBOLD_pred_/d*t* regressor was derived directly from taking the derivative of the predicted BOLD hemodynamic signal—the timing of the visual stimulus paradigm convolved with a canonical double-gamma hemodynamic response function (HRF)—rather than from the measured BOLD signal to improve generalizability. Using dBOLD_pred_/d*t* and its own temporal derivative (to account for slight temporal offsets) as the two regressors, GLM analysis was performed on flow data of all three velocity-encoding directions separately using FSL FEAT^62^. Cardiac-gated CSF flow was generated through retrospective cardiac gating based on voxel-wise timing and alignment with externally-recorded cardiac physiological signals. No temporal filtering was applied for cardiac-gated CSF flow analysis. For BOLD fMRI processing, registered magnitude images were either echo-averaged or fitted to generate T_2_* maps, which were then used to estimate task-related activation (e.g., t-score) using a standard GLM approach using FSL FEAT^62^. No spatial smoothing was applied to the magnitude data. Finally, parameter maps for CSF (velocity, t-score) and BOLD (t-score) were registered to the T_1_-weighted anatomical images for visualization and ROI-based analyses.

### Data analysis and visualization

We used FreeSurfer^63,64^ and its cytoarchitectonic *V1_exvivo*.*label* to automatically define the primary visual cortex (V1) ROI. The V1 subarachnoid space ROI was generated by extending the cortical gray matter ROI outward from the pial surface reconstruction into the adjacent CSF space by 60% of the local cortical thickness using ‘mri_label2vol’ function. V1 cortical gray matter volume and V1 CSF volume were computed based on the FreeSurfer segmentation for correlation analyses with CSF flow. For flow analyses performed within the V1 ROI, the mean value of the 2,000 CSF voxels with the highest flow *t*-scores within the V1 subarachnoid space was calculated for each subject. Statistical comparisons of velocity ranges (**Fig. 2d–g**) were performed using paired two-sided t-tests. Correlations between CSF flow and relevant factors (**Fig. 5a–d** and **Extended Data Fig. 3**), as well as scan–rescan repeatability (**Extended Data Fig. 2a–d**), were evaluated using Pearson’s correlation coefficients (*r*) computed in MATLAB (MathWorks) with the ‘corrcoef’ function. Velocity range was calculated as the difference between the maximum and minimum velocities (can be positive or negative) across frames within the task-locked or cardiac-locked period, and used as a measure of the amplitude of the flow response. The velocity ranges along the three velocity-encoding directions were then combined using root-sum-of-squares to generate the overall velocity range of each voxel, which was used in the ROI analyses.

For visualization, CSF parameter maps (velocity and t-score) and BOLD t-score maps were overlaid on T_1_-weighted anatomical images and displayed using FSLeyes^62^. Three-dimensional flow vector field maps were generated by combining flow measurements along all three directions and visualized in ParaView (Kitware, Inc.) using the “Glyph” representation with “arrow” geometry to depict voxel-wise flow vectors, where vector orientation indicates flow direction, vector length is normalized, and color encodes the combined velocity amplitude.

## ACKNOWLEDGMENTS

This work was supported by the National Institutes of Health (grants R00-AG083056, U24-NS129893, R01-EB036507, R01-AT011429, U19-NS128613, R01-AG070135, R01-EB019437, P41-EB030006, S10-OD023637).

## Author contributions

Conceptualization: ZD, FW; Methodology: ZD, FW; Investigation: ZD, FW, LDL, JRP; Supervision: ZD, FW, LDL, JRP, LLW, BRR; Writing—original draft: ZD, FW; Writing—review & editing: ZD, FW, LDL, JRP, LLW, BRR

## Extended Data

**Extended Data Fig. 1.**
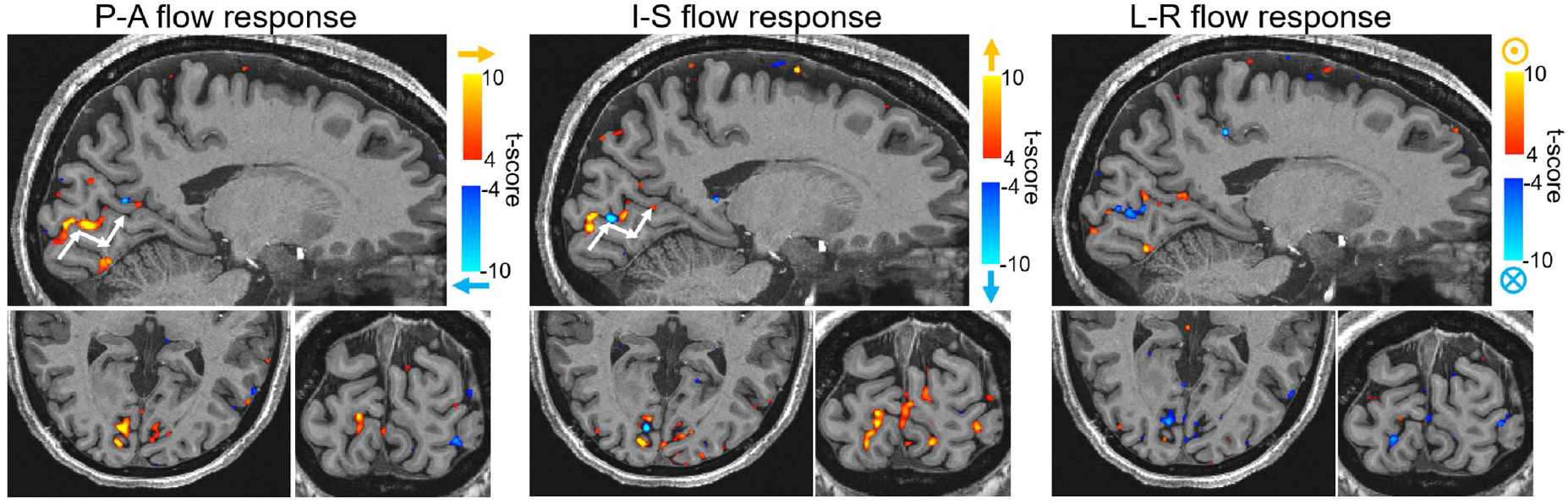
Three-directional flow t-score maps from an example individual. T-score maps for P–A, I–S and L–R directed flow presented in three orthogonal views. White arrows indicate subarachnoid CSF flow directions following the local curvature of the subarachnoid space surrounding the visual cortex.

**Extended Data Fig. 2.**
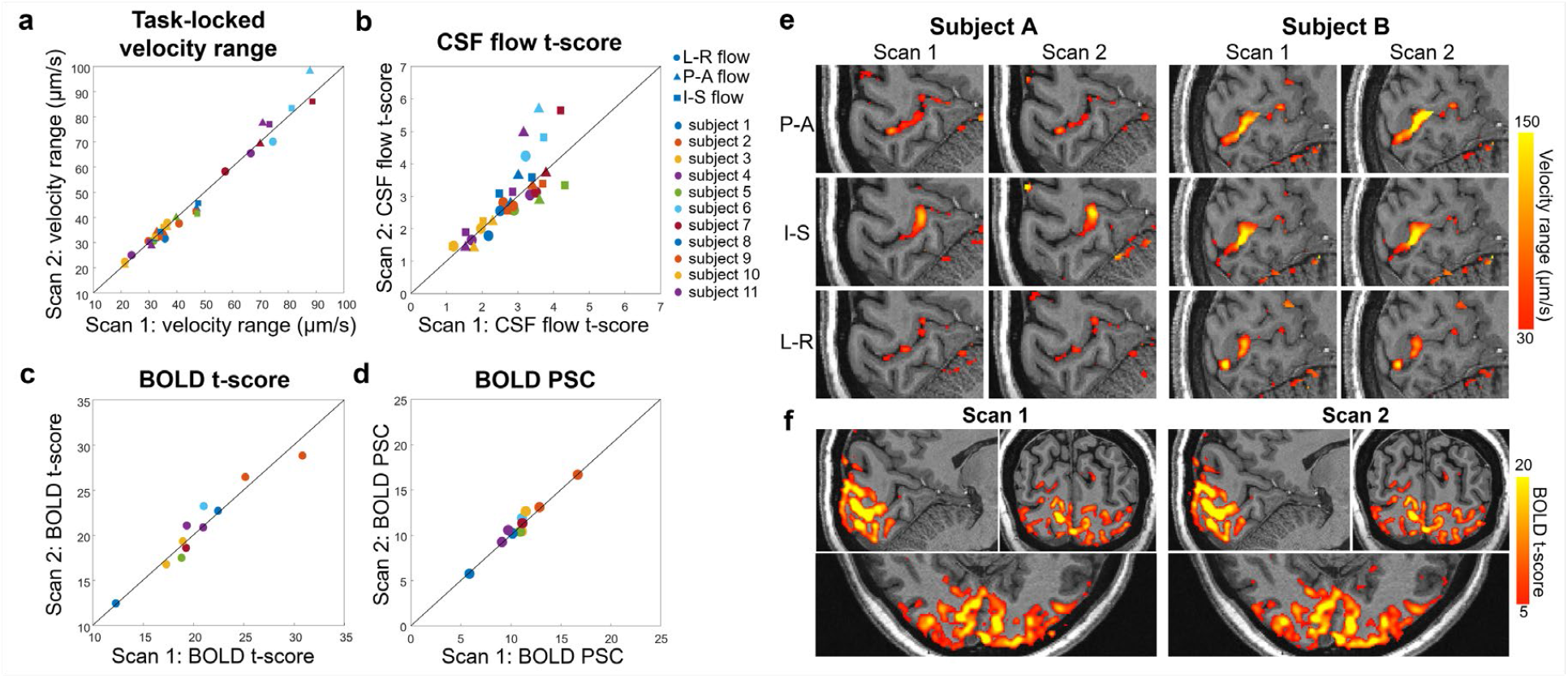
Scan-rescan repeatability of task-evoked CSF flow and BOLD hemodynamic response. **a**, Scatter plot of scan-rescan task-locked velocity ranges (averaged within V1 subarachnoid space) across all participants and all three flow-encoding directions. Different colors denote individual subjects, and different shapes denote flow-encoding directions. The identity line is shown in black. **b**, Scatter plot of scan-rescan CSF flow t-scores (averaged within V1 subarachnoid space) across all participants and all three flow-encoding directions. **c**, Scatter plot of scan-rescan BOLD t-scores (averaged within V1) across all participants. **d**, Scatter plot of scan-rescan BOLD percent signal changes (PSCs) (averaged within V1) across all participants. **e**, Scan-rescan task-locked velocity range maps along the three flow-encoding directions from two representative subjects. **f**, Scan-rescan BOLD t-score maps from a representative subject.

**Extended Data Fig. 3.**
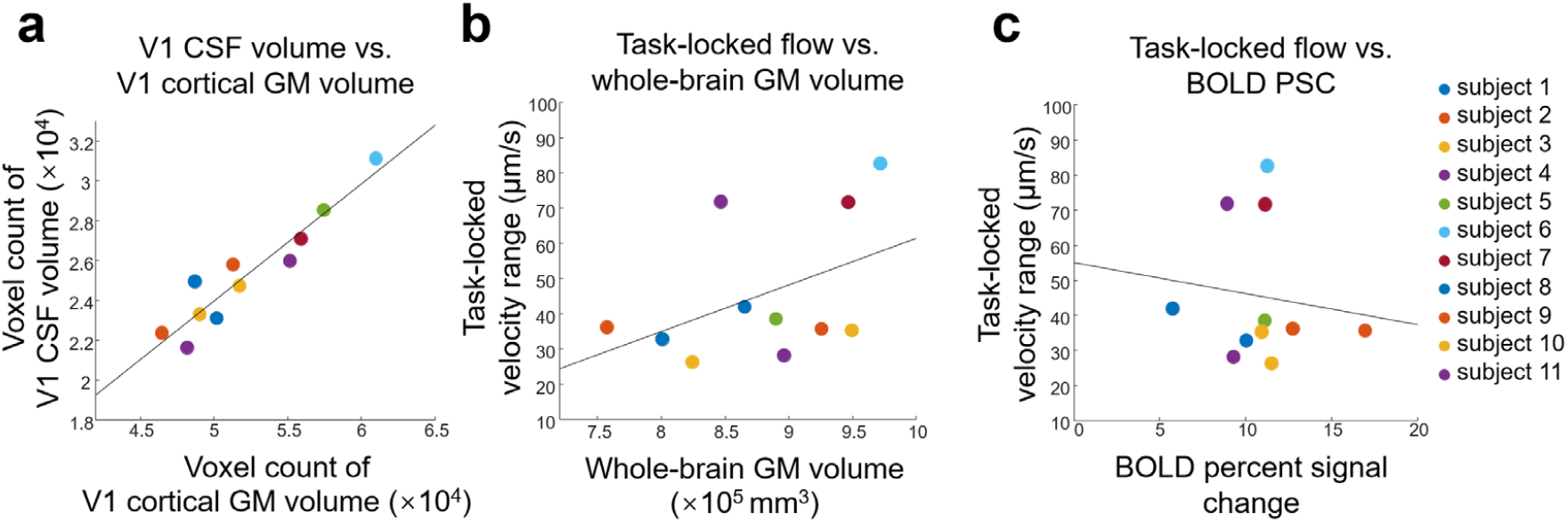
Scatter plots of (**a**) V1 CSF volume versus V1 cortical gray matter volume, (**b**) task-locked velocity range averaged in V1 subarachnoid space versus whole-brain gray matter volume, and (**c**) task-locked velocity range averaged in V1 subarachnoid space versus BOLD percent signal change.

## Supplementary Information

**Supplementary Video 1**. Movie showing visual-task-locked CSF flow velocity and directionality changes along the three velocity-encoding directions across the stimulus cycle.

**Supplementary Video 2**. Movie visualizing CSF flow vector fields within the subarachnoid space surrounding the visual cortex. Blue meshes denote the subarachnoid space, and a midsagittal T_1_-weighted anatomical image is shown for reference. Voxel-wise vector orientation indicates flow direction, vector length is normalized, and color encodes the velocity magnitude.

**Supplementary Video 3**. Movie showing the spatiotemporal evolution of visual-task-locked CSF flow vector fields across the stimulus cycle surrounding the visual cortex, highlighting oscillatory changes in flow velocity and direction, as well as coherent flow patterns following the local curvature of the subarachnoid space. Voxel-wise vector orientation indicates flow direction, vector length is normalized, and color encodes the velocity magnitude.

