## Supplementary figures and images for "Neural activity drives directional subarachnoid cerebrospinal fluid flow in the human brain"

### Supplementary Video 1

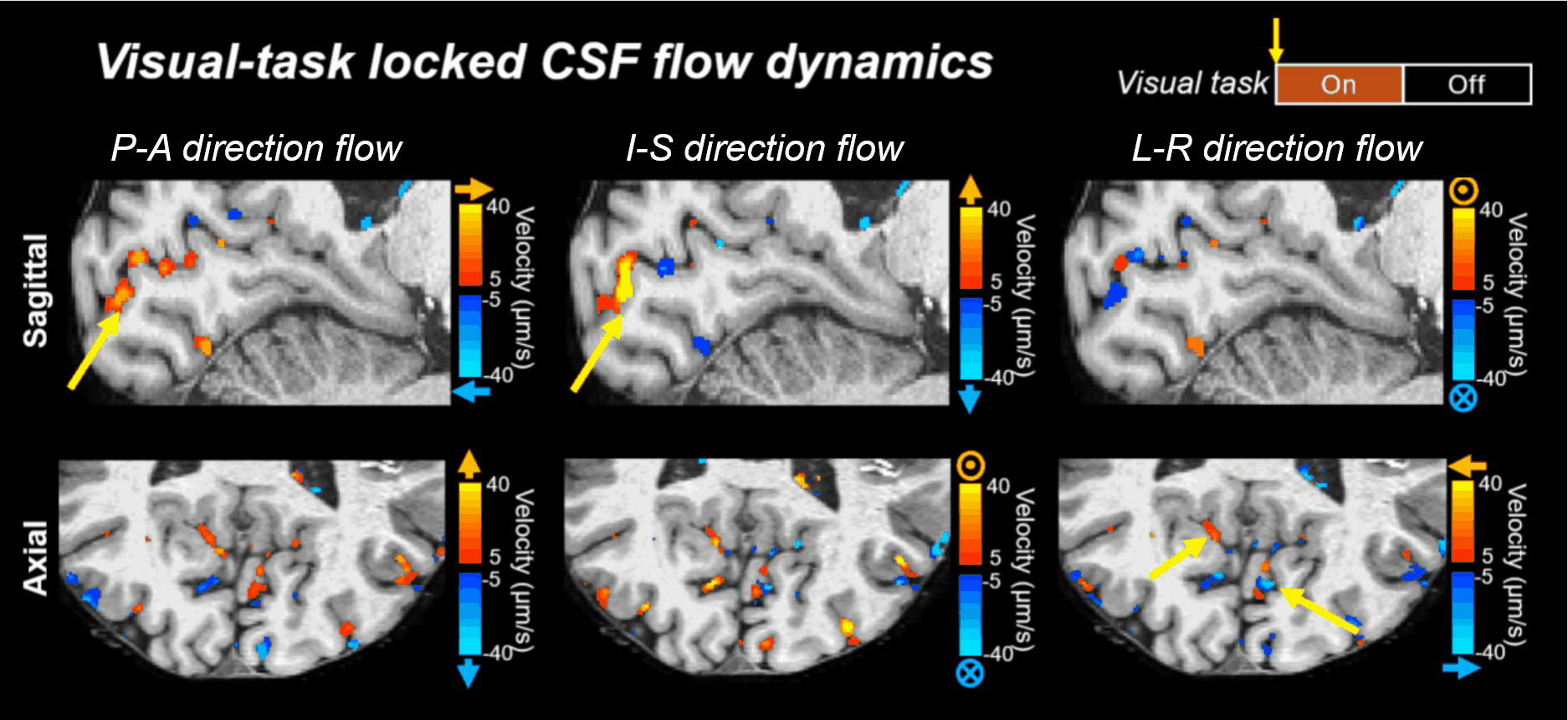

### Supplementary Video 2

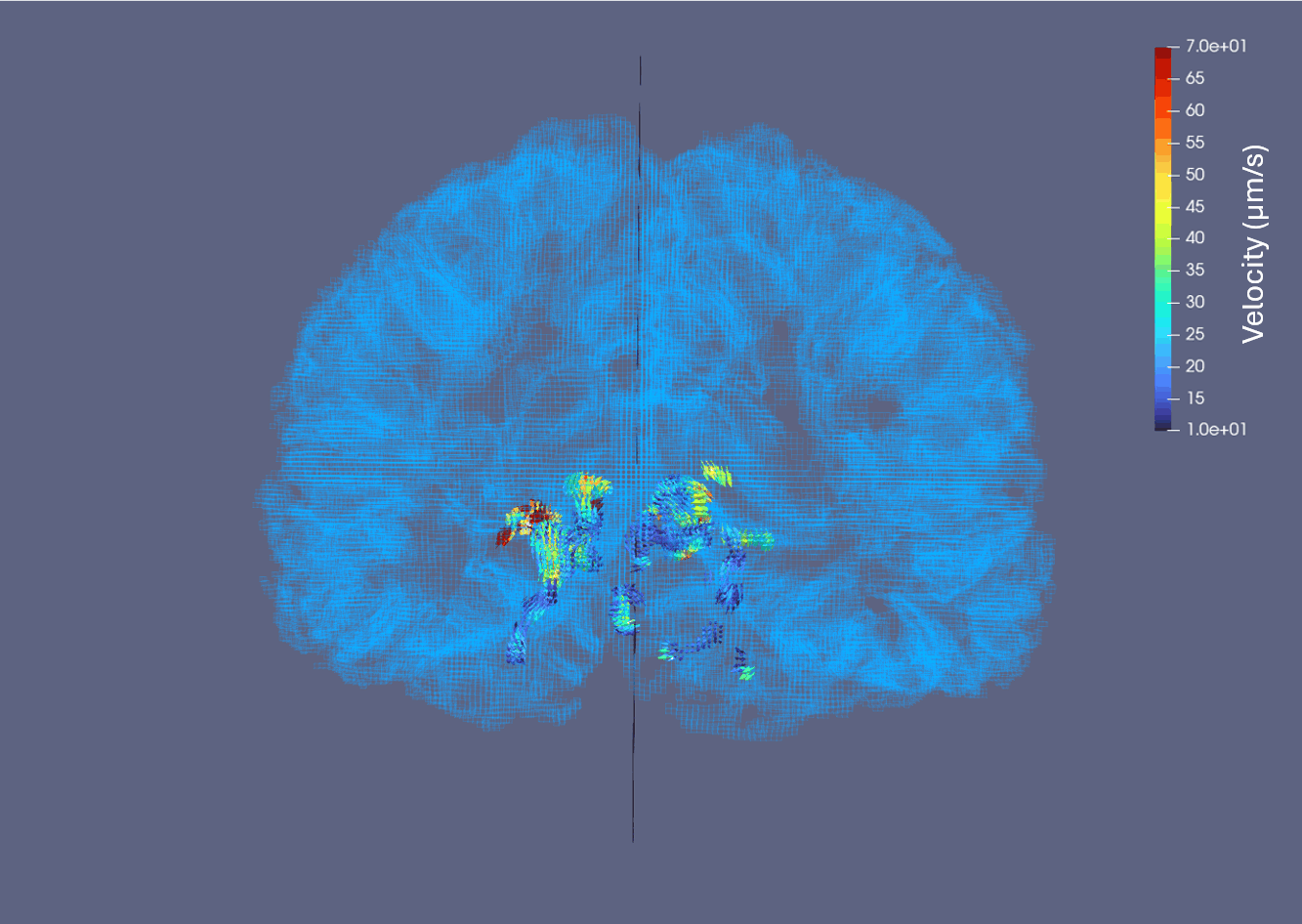

### Supplementary Video 3

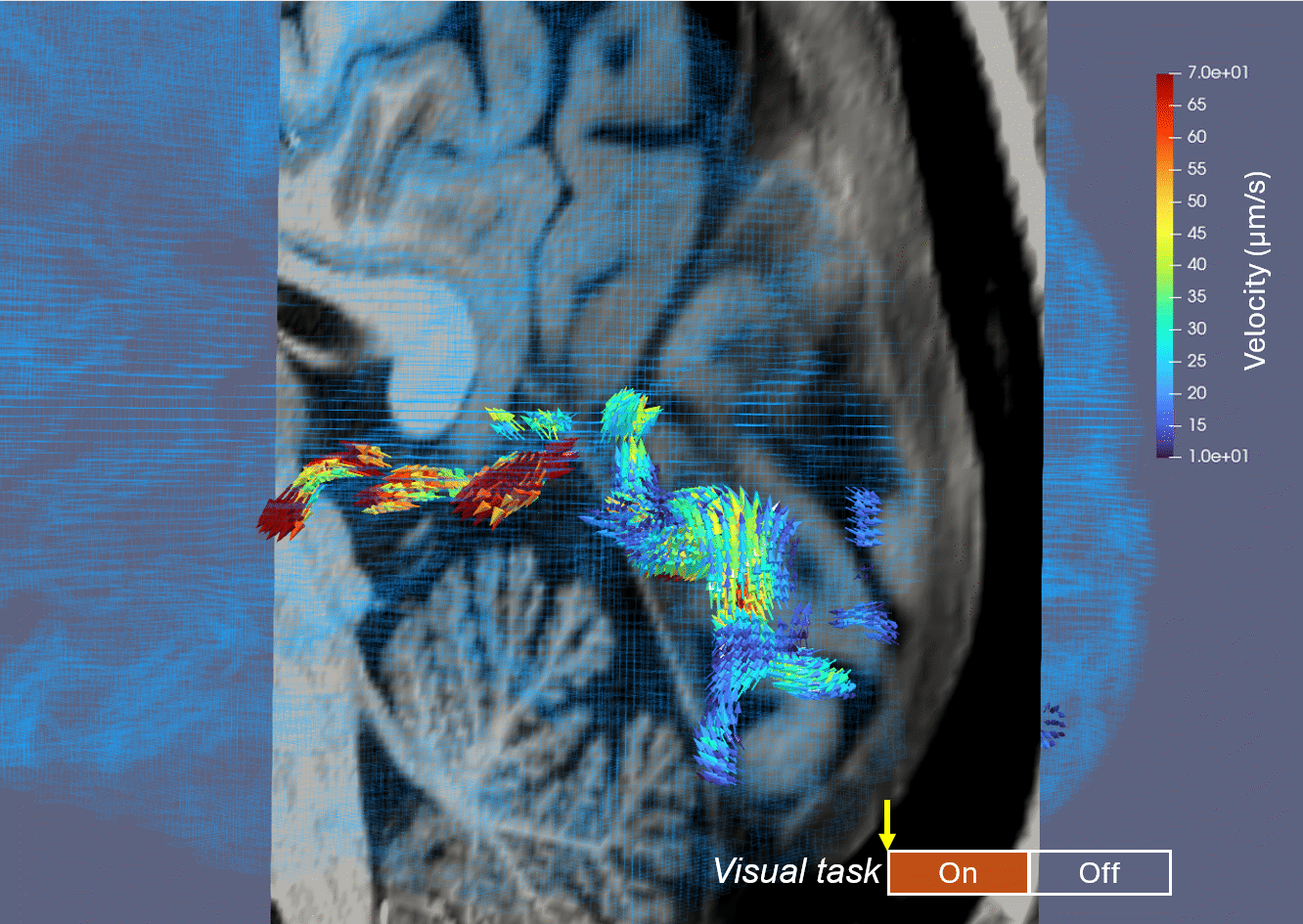
